# Interferon signaling is enhanced by ATR inhibition in glioblastoma cells irradiated with X-rays, protons or carbon ions

**DOI:** 10.1101/2024.06.12.598643

**Authors:** Gro Elise Rødland, Mihaela Temelie, Ana Maria Serban, Adrian Eek Mariampillai, Nina Frederike Jeppesen Edin, Eirik Malinen, Antoine Gilbert, François Chevalier, Diana I. Savu, Randi G. Syljuåsen

**Affiliations:** Department of Radiation Biology, Institute for Cancer Research, Norwegian Radium Hospital, Oslo University Hospital, 0379 Oslo, Norway; Department of Life and Environmental Physics, Horia Hulubei National Institute of Physics and Nuclear Engineering, 077125 Magurele, Romania; Department of Physics, University of Oslo, 0371 Oslo, Norway; UMR6252 CIMAP, Team Applications in Radiobiology with Accelerated Ions, CEA-CNRS-ENSICAEN-Université de Caen Normandie, 14000 Caen, France

## Abstract

**Background and purpose:** Interferon signaling plays an important role in antitumor immune responses. Inhibitors of the DNA damage response, such as ATR inhibitors, can increase interferon signaling upon conventional radiotherapy with X-rays. However, whether such inhibitors also increase interferon (IFN) signaling after high linear energy transfer (LET) particle irradiation is not known.

**Materials and methods:** Human glioblastoma U-251 and T98G cells were treated with X-rays, protons (linear energy transfer (LET): 7 and 38 keV/μm) and carbon ions (LET: 28 and 73 keV/μm), with and without ATR inhibitor (VE822) or ATM inhibitor (AZD1390). DNA damage signaling and cell cycle distribution were assayed by immunoblotting and flow cytometry, and radiosensitivity by clonogenic survival. IFN-β secretion was measured by ELISA and STAT1 activation by immunoblotting.

**Results:** High-LET protons and carbon ions caused stronger activation of the DNA damage response compared to low-LET protons andX-rays at similar radiation dose. G2 checkpoint arrest was abrogated by the ATR inhibitor and prolonged by the ATM inhibitor after all radiation types. The inhibitors increased radiosensitivity, as measured after X- and carbon-ion-irradiation. ATR inhibition increased IFN signaling after both low-LET and high-LET irradiation in both cell lines. In T98G, IFN signaling was also enhanced by ATM inhibition. Notably, T98G cells secreted markedly more IFN-β when the inhibitors were combined with high-LET compared to low-LET irradiation.

**Conclusion:** Our results show that ATR inhibition can increase IFN signaling after both X-, proton- and carbon-ion-irradiation. Additionally, IFN induction is strongly dependent on LET in one of the tested cell lines.

## Introduction

Glioblastoma (GBM) is the most common and deadliest type of brain cancer. Radiotherapy is part of the standard treatment, but treatment efficacy is limited by tumor radioresistance and damage to the normal brain [1]. Particle irradiation with carbon ions or protons may potentially improve GBM treatment [2, 3]. The advantageous depth dose distribution of particle beams allows sparing of normal tissue [4, 5]. Furthermore, carbon ions or protons with high linear energy transfer (LET) induce clustered DNA damage, which can be harder to repair and thus more potent in eliminating tumor cells [2]. To further enhance the tumor cell killing, radiotherapy may be combined with inhibitors of cell cycle checkpoints and DNA repair pathways [6, 7]. It was previously thought that such inhibitors would not sensitize to high-LET irradiation, based on an assumption that the clustered DNA damage would be mostly irreparable [8]. However, more recent studies indicate that DNA repair is also important upon high-LET irradiation [9-11]. Radiosensitizing effects have for example been observed with ATR and PARP inhibitors in combination with carbon ion irradiation [12-16].

Interestingly, combining radiotherapy with DNA damage response inhibitors may also trigger antitumor immune effects. For instance, abrogation of the radiation-induced G2 checkpoint by ATR inhibition results in increased micronucleus formation that can induce a type 1 interferon (IFN) response in various cell types. The micronuclei are prone to rupture, and DNA from ruptured micronuclei can be recognized by the cytosolic DNA sensor cGAS, leading to type 1 interferon secretion via the cGAS-STING-TBK1-IRF3 pathway. Alternatively, increased IFN production may be triggered by cytosolic RNA via the RNA sensor RIG-I [17, 18]. Inhibiting ATR or ATM in combination with irradiation also promotes antitumor immune responses in mouse tumor models [19-22]. Moreover, ongoing clinical trials are investigating combinations of ATR inhibitors and immunotherapy, with and without radiotherapy [23].

While previous preclinical studies have shown enhanced antitumor immune signaling upon combination of DNA damage response inhibitors and X-irradiation, it is not clear whether such inhibitors also increase the immune signaling when combined with proton or carbon ion irradiation. Furthermore, little is known about interferon signaling after irradiation and DNA damage response inhibition in GBM. Here we show that ATR inhibition can abrogate the G2 checkpoint and increase type 1 IFN signaling in two GBM cell lines after X-, proton and carbon-ion-irradiation. In addition, ATM inhibition increases IFN signaling in one of the cell lines, despite a lack of G2 checkpoint abrogation. Our results suggest that DNA damage response inhibitors could be useful in combination with proton or carbon ion therapy to increase tumor cell radiosensitivity and type 1 IFN responses.

## Materials and Methods

### Cell culture

Human glioblastoma cell lines T98G and U-251 were cultured in Dulbecco’s modified Eagle’s medium supplemented with 10% fetal bovine serum (Biowest) and 1% penicillin/streptomycin solution (10000 U/mL) (ThermoFisher Scientific), at 37°C in a humidified atmosphere with 5% CO_2_. T98G was kindly provided by Dr. Theodossis Theodossiou and U-251 was purchased from CLS Cell Line Servic (Eppelheim, Germany). Both cell lines were verified by short tandem repeat analysis (Eurofins).

### Drug treatment

Inhibitors of ATM (AZD1390) and ATR (VE-822/berzosertib), both from Selleck Chemicals, were given to cells 15-30 min prior to irradiation for X-ray and carbon ion experiments, or immediately after irradiation for proton experiments. In clonogenic survival assays the inhibitors were removed after 24 hours of exposure, otherwise they were present until the end of the experiment.

### Cell irradiation

Proton irradiation was performed at the Oslo cyclotron laboratory (University of Oslo) with an MC-35 cyclotron (Scanditronix) with a beam energy of 15.5 MeV. The beamline and setup has been described previously [24, 25]. The cells were irradiated in two positions; in front of the Bragg peak and at the distal end. The dose rate was the same for both positions. The medium was removed during irradiation and fresh medium was added immediately after irradiation. Carbon ion irradiation was performed using the IRABAT 95 MeV beam line at the GANIL facility (Caen, France), with further details described in [26]. X-rays (160 kV, 6.3 mA, Faxitron) were delivered at 1 Gy/min (filtration 0.8 mm Be + 0.5 mm Cu).

The LET values for carbon ions were estimated at 28 and 73 keV/μm and for protons 7 and 38 keV/μm. In our study we have termed the lowest LET value for each particle type “low-LET” with the purpose of separately comparing the two LET values within each radiation modality. The LET for 160 kVp X-rays is ∼4 keV/μm [27].

### Flow cytometry

Cells were fixed in 70% ice cold ethanol. To eliminate sample-to-sample variation, we added an aliquot of barcoded reference cells stained with Alexa Fluor 647 Succinimidyl Ester (0.02 μg/μL) (ThermoFisher Scientific) to all samples. Samples were stained with mouse anti-γH2AX (Ser139), clone JBW301 (Millipore), conjugated to FITC (1:1000) or not (1:500), the latter followed by Alexa Fluor 488 anti-mouse IgG (1:500, Molecular Probes). DNA was stained with Hoechst 33258 (Sigma-Aldrich). Samples were analysed on an LSR II flow cytometer (BD Life Sciences) or on a CytoFLEX (V5-B4-R3) (Beckman Coulter). Results were analysed in FlowJo v.10.6.1 software (BD Life Sciences), using the Watson Pragmatic algorithm for cell cycle analysis. Instrument service on the LSR II was provided by the Flow Cytometry Core Facility (OUS).

### Clonogenic survival assay

Cells were seeded in triplicate 6 cm dishes (500-6000 cells/dish), 18-24 h prior to treatment with X-rays and inhibitors. When colonies appeared (10-14 days), cells were fixed with 70 % ethanol and stained with methylene blue. Colonies with >50 cells were counted. For experiments with carbon ions, the same procedure was followed, except that cells were seeded in T-25 culture flasks at densities of 750-15000 cells/flask.

### Western blotting

Cell lysis and immunoblotting was performed as previously described [28], with the exception that Criterion TGX Stain-free gels (Bio-Rad) were used. Protein concentration was not measured in samples shown in Figure 1. The antibodies used are listed in Table S1. Protein bands were quantified in the ImageLab 4.1 software. A dilution series of one of the samples was included for accurate quantification.

**Figure 1.**
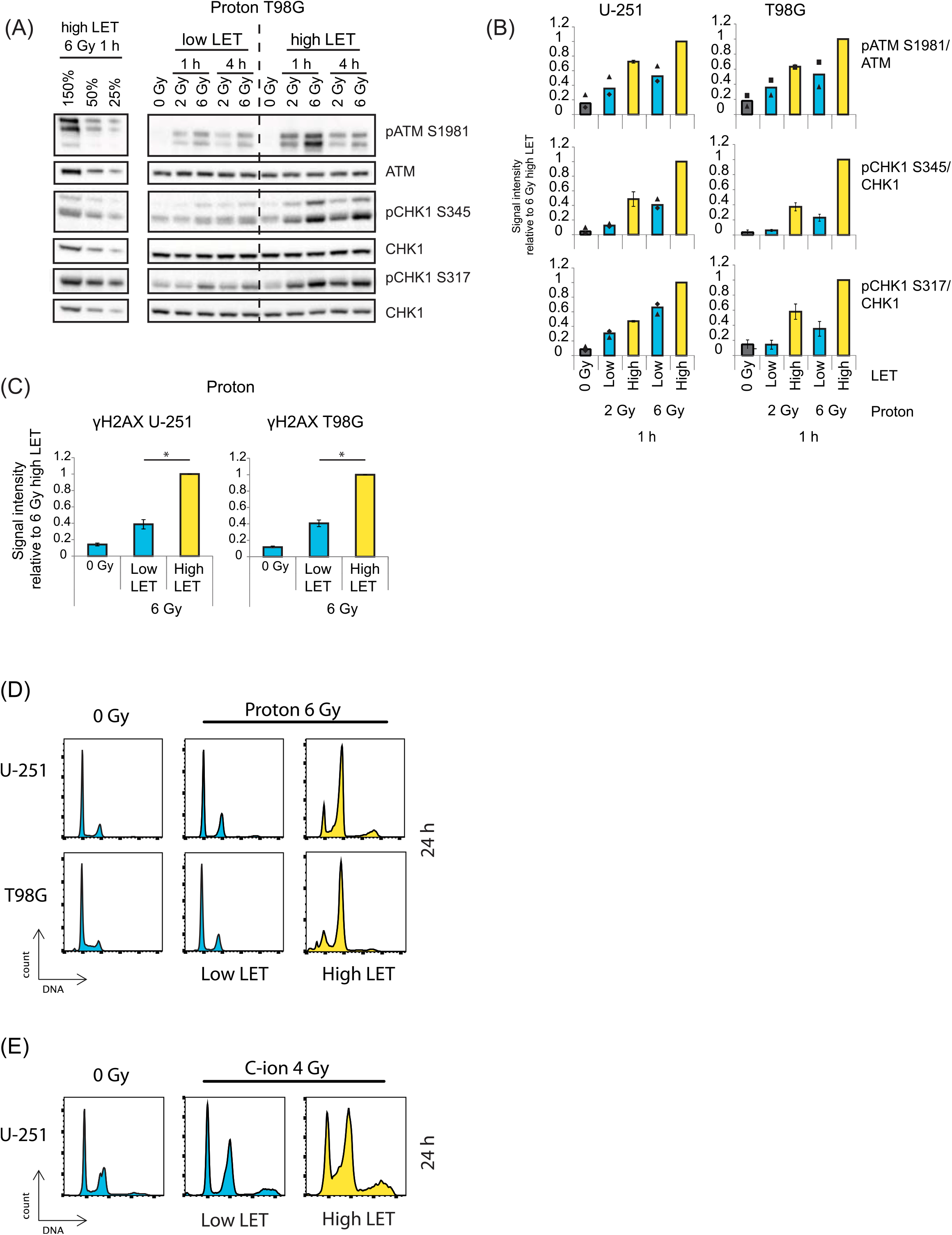
Activation of the DNA damage response is stronger in GBM exposed to high-LET as compared to low-LET irradiation at similar radiation dose. (A) Representative immunoblot from T98G, showing phosphorylated ATM and CHK1 at 1 and 4 h after 2 and 6 Gy of proton irradiation. Left: a dilution curve showing dynamic range of antibodies. (B) Quantification of signal intensity of the indicated markers from two or more experiments similar to that shown in A, for both T98G and U-251. Values are normalized to the 6 Gy high-LET sample. Dots indicate individual experiments. Error bars: SEM (*n* = 3). (C) Quantification of γH2AX intensity as measured by flow cytometry 0.5 h after proton irradiation. Error bars: SEM (*n* = 4). (D) DNA profiles obtained by flow cytometry showing cell cycle distribution at 24 h after exposure to 6 Gy of low-vs. high-LET protons. Upper panels: U-251, lower panels: T98G. (E) Cell cycle distribution of U-251 cells 24 h after irradiation with 4 Gy of carbon ions (C-ions), comparing low and high LET.

### Enzyme-linked immunosorbent assay (ELISA) of secreted IFNβ

Growth medium supernatants were collected from the cell samples. Duplicates of 50 μl of concentrate were subjected to IFN-β measurement by ELISA (Human IFN-beta DuoSET ELISA, R&D Systems), after 20X upconcentration (Amicon Ultracel-10, Merck) of the supernatants as previously described [28]. The protein concentration of remaining adherent cells at time of harvest was measured as described above for western blot samples, and used for normalization of ELISA IFN-β read-outs.

### Statistics

One- or two-sample, two-tailed Student’s *t* test with significance level set to 0.05 was used to obtain *p*-values, indicated with an asterisk for *p* < 0.05 in bar charts.

## Results

To investigate effects of ATM and ATR inhibitors in combination with low- and high-LET irradiation, we first characterized DNA damage signaling after irradiation alone. Immunoblotting on samples harvested at 1 and 4 h after proton irradiation showed that high-LET protons induced more phosphorylation of ATM and the ATR target CHK1 compared to low-LET protons at similar radiation dose (Fig. 1A, 1B). These effects were seen in both T98G and U251 cells. Furthermore, flow cytometry analysis of the DNA-damage marker γH2AX at 0.5 h after irradiation also showed much stronger signals for the high-LET as compared to low-LET protons (Fig 1C). The γH2AX levels in T98G exposed to 6 Gy of high-LET protons were comparable to that observed in cells X-irradiated with 12 Gy (Fig. S1A, S1B). To study potential differences in repair rate between cells exposed to low- and high-LET irradiation we also assessed γH2AX at 24 h. The ratio between the signal at 24 and 0.5 h was similar for cells irradiated with 6 Gy of low-LET protons and X-rays, whereas the ratio for 6 Gy of high-LET protons was comparable to that seen for 15 Gy X-ray (Fig. S1B, right). The high-LET protons thus likely induce DNA damage that is harder to repair.

Since DNA damage signaling leads to activation of cell cycle checkpoints, we also obtained cell cycle profiles at 24h after low- and high-LET proton and carbon ion irradiation. We found that activation of the G2 checkpoint, in both U-251 and T98G, was stronger upon 6 Gy of the high-LET than low-LET proton irradiation, as seen by the presence of more cells in G2/M phase (Fig. 1D). The high-LET carbon ions (73 keV/μm) also showed stronger G2 checkpoint activation compared to the low-LET carbon ions (29 keV/μm) (Fig. 1E). These results are in agreement with more CHK1 phosphorylation observed with high-LET irradiation (Fig 1A, 1B), as the radiation-induced G2 checkpoint is typically dependent on ATR-CHK1 activation [6].

Next, we investigated how the radiation-induced G2 arrest was affected by co-treatment with inhibitors of the DNA damage response proteins ATR and ATM. In line with previous studies [28, 29], the ATR inhibitor abrogated, and the ATM inhibitor prolonged, G2 arrest at 24 h after irradiation with X-rays in both cell lines (Fig. 2A). These effects of the inhibitors were also seen with high-LET carbon ions and protons (T98G) (Fig. 2B, 2C, S2). Moreover, both inhibitors decreased clonogenic survival of U-251 cells when combined with X-rays and both LETs of carbon ions (Fig. 3). These data add to the studies showing that DNA-repair inhibitors can indeed sensitize cells to high-LET particle irradiation [13-15]. However, the sensitizing effect of the inhibitors appeared slightly smaller in combination with the high-LET carbon ions compared to with X-rays and was lower for the ATM inhibitor than the ATR inhibitor (Fig. 3 and Table 1).

**Table 1.** RBE at SF_50_.

|  | Dose (Gy) |  |  | RBE |  |
| --- | --- | --- | --- | --- | --- |
|  | X-ray | C-ion |  | C-ion |  |
|  |  | low | high | low | high |
| - | 3.0 | 1.6 | 1.1 | **1.9** | **2.7** |
| ATMi 10 nM | 1.5 | 0.9 | 0.8 | **1.7** | **2.0** |
| ATRi 100 nM | 2.0 | 1.1 | 0.9 | **1.9** | **2.3** |

RBE values of C-ions with radiation doses of C-ions and X-rays required for killing 50 % of U-251 cells in a clonogenic survival assay.

**Figure 2.**
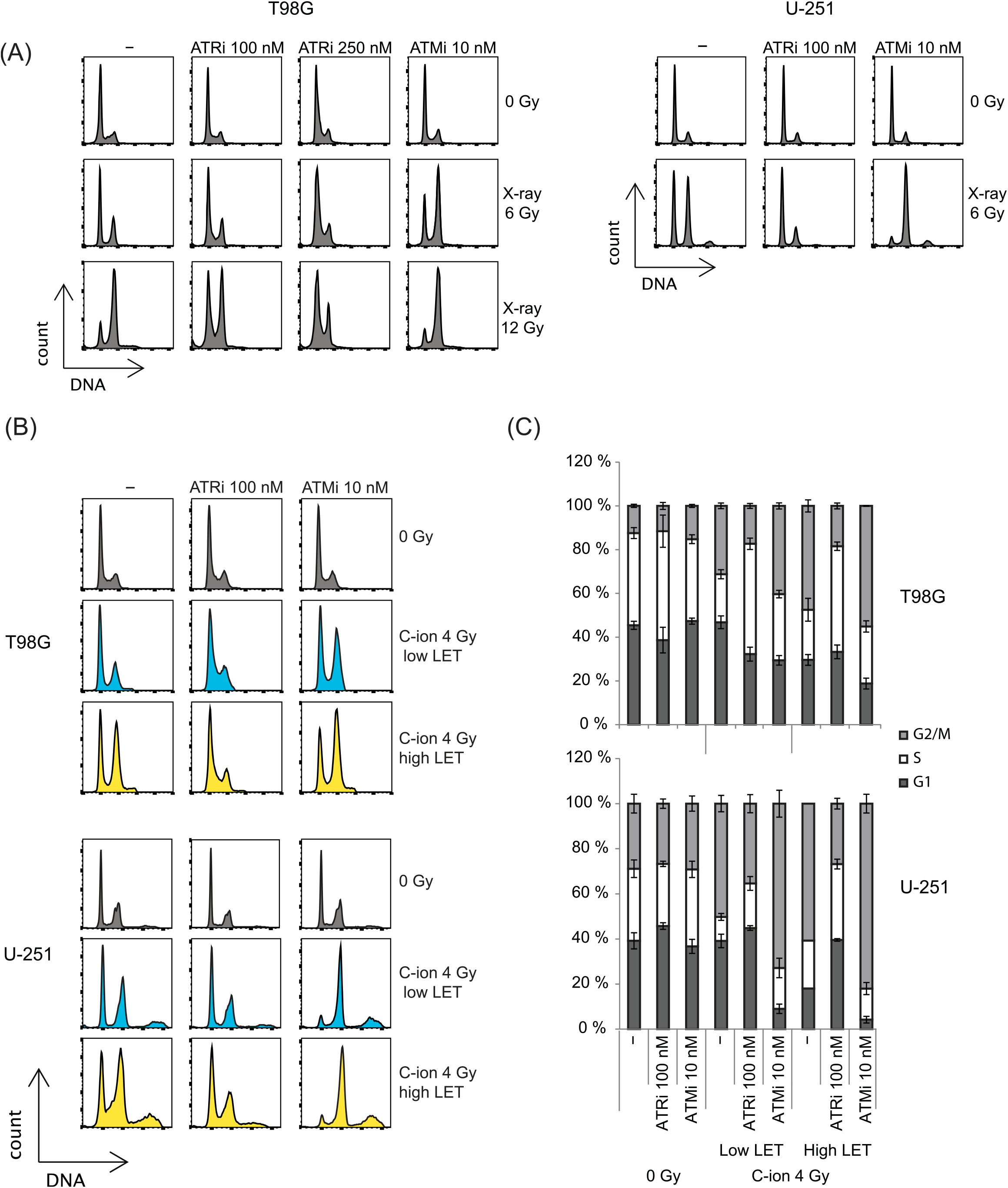
The radiation-induced G2 checkpoint is abrogated by ATR inhibition and prolonged by ATM inhibition. (A) DNA profiles showing cell cycle distribution at 24 h after treatment with ATR and ATM inhibitors in combination with X-irradiation. Left: T98G, right: U-251. (B) DNA profiles at 24 h after treatment with the inhibitors and carbon ions, comparing low and high LET. (C) Quantification from cell cycle analysis performed on data as in B. Error bars: SEM (*n* = 3).

**Figure 3.**
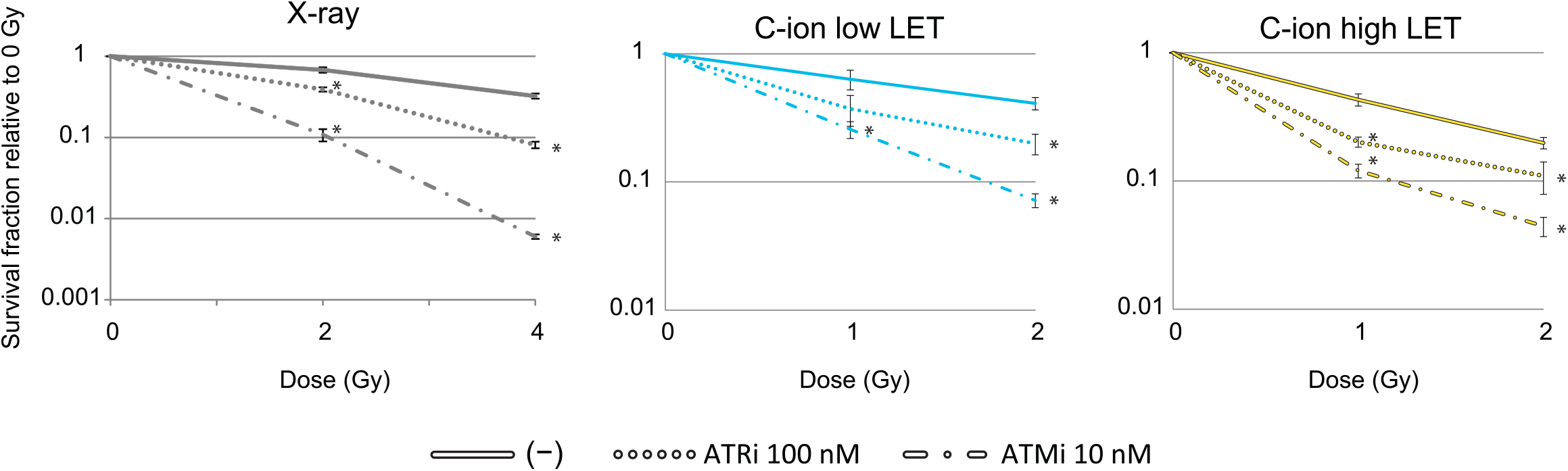
ATM and ATR inhibitors enhance radiosensitivity after both low and high-LET irradiation. Clonogenic survival of U-251 cells irradiated with 2 and 4 Gy of X-ray (left panel) or 1 and 2 Gy of carbon ions (middle and right panels) in combination with ATR and ATM inhibitors. Survival fractions relative to non-irradiated samples are plotted. The average survival fractions after treatment with the inhibitors alone were 0.6 (ATRi) and 0.8 (ATMi) for the experiments with X-rays, and 0.7 (ATRi) and 0.9 (ATMi) for the experiments with carbon ions. Error bars: SEM (*n* = ≥3).

Since abrogation of the radiation-induced G2 cell cycle checkpoint by ATR inhibition is known to induce a type 1 interferon response upon X-irradiation, we asked whether similar responses would be induced after high-LET irradiation. We first examined phosphorylation of STAT1 (phospho-STAT1), a commonly used surrogate marker for extracellular type 1 IFN [30], by immunoblotting of U-251 cells three days after X- and carbon-ion-irradiation in combination with inhibitor treatment. The time point was chosen as previous studies have shown increased IFN signaling three days after ATR inhibition and X-irradiation in various cancer cell lines [18, 28]. Phospho-STAT1 was increased by ATR inhibition, but very little by ATM inhibition, after 4-6 Gy of X-irradiation (Fig. 4A). Similar results were obtained in cells irradiated with 4 Gy of carbon ions, with little difference between the low- and high-LET (Fig. 4B). The amount of IFN-β in the growth medium was also directly measured by ELISA in U-251 cells treated with X-irradiation and inhibitors. The total levels varied between experiments (Fig. S3A), but the highest levels were detected after co-treatment with ATR inhibitor and X-rays, and overall correlated well with phospho-STAT1 signals measured in corresponding cell lysates (Fig. 4C). Notably, the observed phospho-STAT1 levels of samples irradiated with carbon ions would not correspond to higher IFN-β levels compared to X-irradiated samples (Fig. 4C, red color).

**Figure 4.**
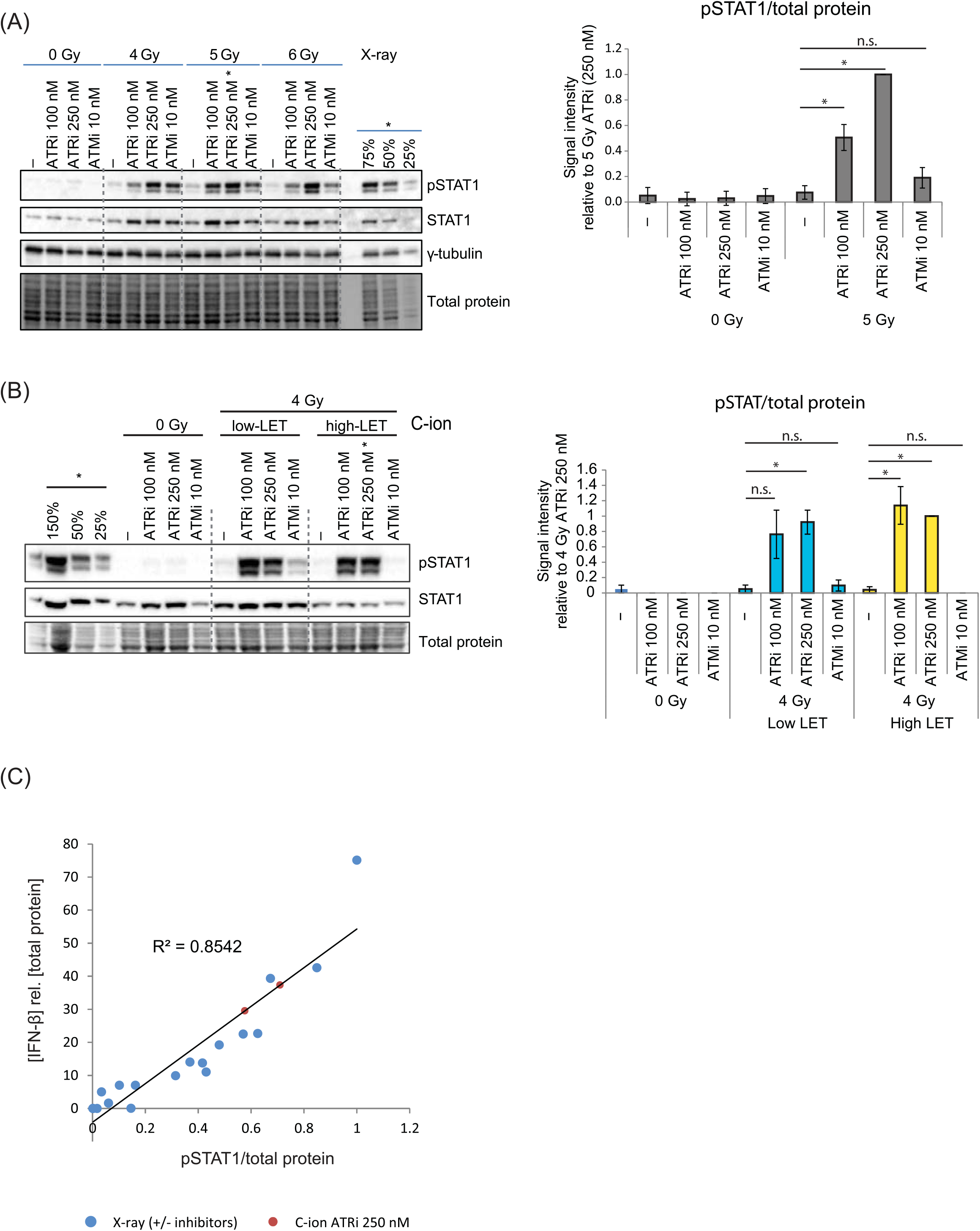
ATR inhibition increases type 1 IFN signaling after both low-LET and high-LET irradiation in U-251 cells. (A) Left: Representative immunoblot showing phosphorylated STAT1 (pSTAT1) in U-251 cells at 72 h after treatment with 4, 5 and 6 Gy of X-ray in combination with indicated inhibitors. Asterisk indicates a dilution of the sample treated with 5 Gy and 250nM ATRi. Right: Quantification of pSTAT1 relative to total protein, normalized to the sample treated with 5 Gy in combination with 250 nM of ATRi, Error bars: SEM (*n* = 7). (B) Left: Representative immunoblot showing pSTAT1 in U-251 cells at 72 h after treatment with 4 Gy of low- and high-LET carbon ion irradiation in combination with indicated inhibitors. Asterisk indicates a dilution of the sample treated with 4 Gy high LET and 250nM ATRi. Right: Quantification of pSTAT1 relative to total protein, normalized to the sample treated with 4 Gy high LET in combination with 250 nM of ATRi, Error bars: SEM (*n* = 3). (C) Correlation analysis between pSTAT1 and IFN-β secretion in X-irradiated samples. Signal intensity of pSTAT1 as measured by immunoblotting on samples as in A (relative to 6 Gy ATRi 250 nM) are plotted against levels of secreted IFN-β measured in growth medium from the same samples. The IFN-β values have been normalized to the amount of protein of the adherent cells at time of harvest. The red dots show pSTAT1 signals from carbon ion irradiated samples that have been analyzed in the same immunoblot as the X-irradiated samples, in order to estimate the IFN-β level upon co-treatment with C-ions and ATRi. (n.s.: not significant).

Phospho-STAT1 levels were also detected by immunoblotting in T98G cells treated with combinations of inhibitors and X-ray, proton or carbon ion irradiation. For all radiation modalities, an increase in phospho-STAT1 signal was observed for both inhibitors as compared to irradiation alone (Fig. 5A, S3B). However, the increase was not particularly strong and appeared similar for low- and high-LET particles. Moreover, the signal intensity did not correlate well with IFN-β secretion measured in the cell growth medium in X-irradiated samples (Fig. S3C). This could be due to partly defective IFN receptor signaling as has been reported for other cancer cell lines [28]. Interestingly, IFN-β levels in the growth medium of T98G cells exposed to high-LET particle irradiation in combination with the inhibitors were clearly higher than that measured in growth medium of cells exposed to low-LET particles or X-rays (Fig. 5B). This was found also when compared to very high doses of X-rays (30 Gy). IFN-β levels were high for both LETs of carbon ions co-treated with ATR inhibitor, whereas only the high-LET irradiation produced such high levels for protons.

**Figure 5.**
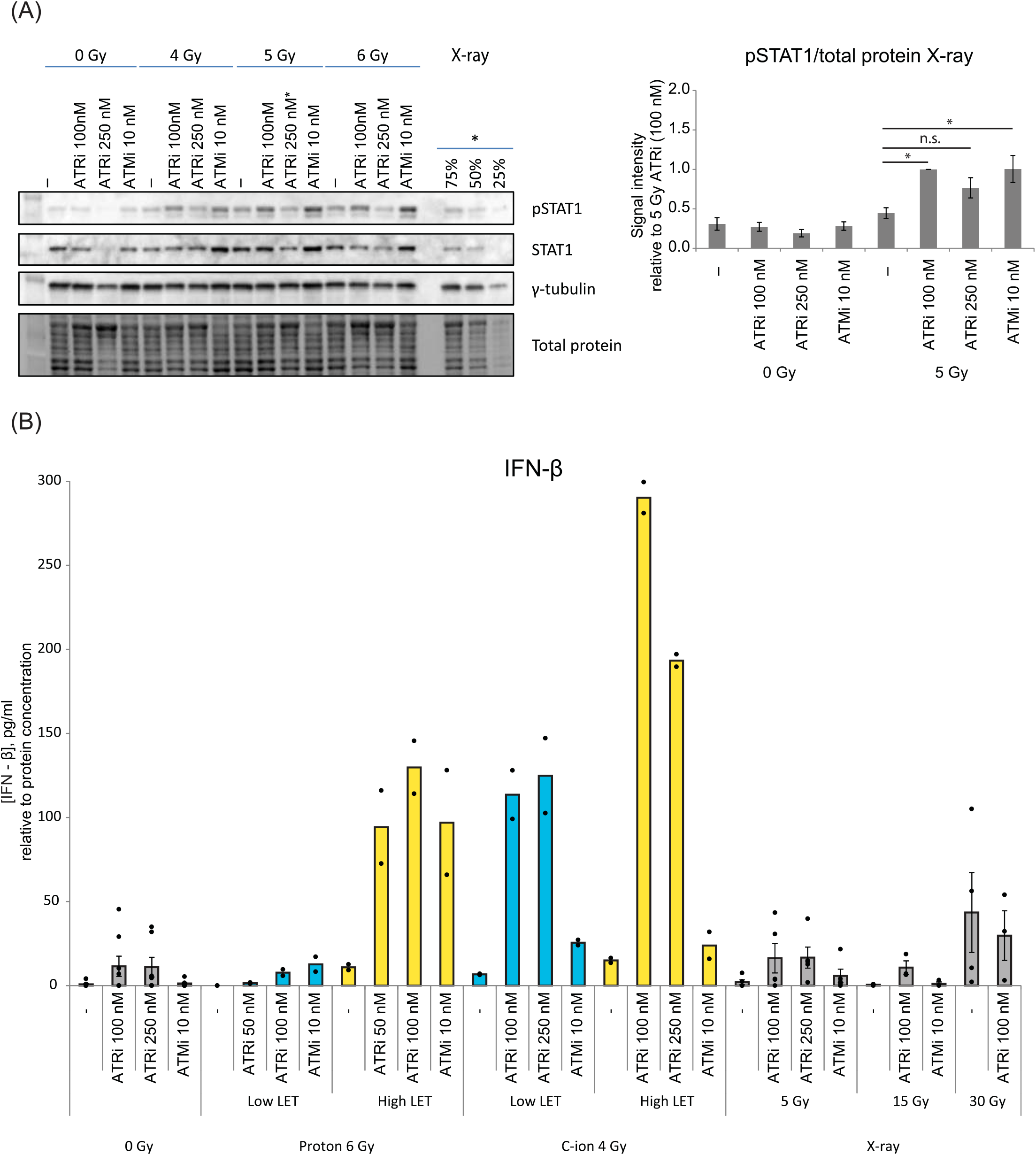
IFN signaling is heavily induced in T98G cells treated with high-LET irradiation combined with ATR inhibitor. (A) Analysis of pSTAT1 in T98G cells, similarly as in Fig 4A except that values are normalized to the sample treated with 5 Gy and 100 nM ATRi. Error bars: SEM (*n* ≥ 5). (B) Levels of secreted IFN-β measured by ELISA relative to the amount of protein of the adherent cells at time of harvest, comparing all radiation modalities. Proton and carbon ion irradiated samples are from the same experiments as in S3B. For X-ray irradiated samples *n* ≥ 3. Data from individual experiments are displayed as dots.

However, as previously mentioned, the low LET of carbon ions is comparable to the high LET of protons. Also noteworthy is that in combination with proton irradiation, inhibition of ATM leads to an IFN-β secretion level similar to that seen after ATR inhibition (Fig. 5B). This was not the case for samples irradiated with carbon ions. Taken together, these results show that ATR inhibition increases IFN secretion after irradiation with X-rays, protons or carbon ions in both cell lines, and in T98G this is also highly LET-dependent. In addition, ATM inhibition can increase radiation induced IFN-β secretion in T98G.

## Discussion

Radiotherapy with high energy X-rays is a cornerstone of cancer treatment. Recently, the interest in understanding radiation-induced immune effects has escalated, with the goal to optimize combination treatments with immunotherapy. Inhibitors of DNA repair may counteract tumor radioresistance, and may in addition enhance the antitumor immune effects. Particle radiotherapy with protons or carbon ions is currently being expanded in many countries, and increased immune signaling has also been observed in response to these radiation modalities [31, 32]. However, knowledge about how DNA repair inhibitors modulate antitumor immune effects after particle irradiation has been lacking. To our knowledge, our study is the first to report effects on the IFN response after treatment with ATR and ATM inhibitors in combination with high-LET particle irradiation.

Notably, these inhibitors may be even more suitable for combinations with particles as opposed to conventional X-ray radiotherapy. The radiosensitizing effects of the inhibitors on the surrounding normal tissue, including immune cells, will likely be reduced, due to the beneficial depth dose distribution of particle radiation. We thus expect improved tumor-selective radiosensitizing effects and possibly improved antitumor immune response with particle irradiation. In support of the latter, recent clinical reports suggest that proton and carbon ion radiotherapy induce less lymphopenia compared to classical radiotherapy [33, 34]. Our data suggest that a further benefit from combining ATR and ATM inhibitors with particle irradiation may be obtained through activation of the innate immune response within the tumor cells themselves.

In U-251 cells the radiation-induced IFN-β signaling correlated with abrogation of the G2 checkpoint. Cells treated with the ATR inhibitor were not arrested and showed increased IFN-β signaling. The ATM inhibitor, by contrast, prolonged the G2 arrest and gave little increase in IFN signaling. In T98G cells, both inhibitors induced IFN-β signaling, even though their effect on the G2 checkpoint was similar to that observed in U-251. Cytosolic exposure of DNA from ruptured micronuclei is most likely a main cause for increased IFN-β production after ATR inhibition in both cell lines, as previously reported for combination with X-irradiation in other cell types [28, 35]. However, other mechanisms are likely at play after ATM inhibition, as the G2 arrest is prolonged. In the absence of irradiation, ATM inhibition causes leakage of mitochondrial DNA into the cytosol [36]. This response may be enhanced by radiation-induced mitochondrial damage. Furthermore, a more direct effect of ATM inhibition on signal transduction has been shown, where reduced ATM activity causes elevated SRC phosphorylation leading to TBK1 activation [22].

Interestingly, the levels of secreted IFN-β from T98G were much higher when the inhibitors were combined with high-LET than low-LET irradiation. The mechanism underlying this effect is not known and would be interesting to explore in future studies. Pointing towards possible mechanisms, studies have shown that more micronuclei are formed in cells exposed to high-LET than low-LET irradiation [13, 37, 38], likely due to induction of more complex DNA damage. Furthermore, in addition to cytosolic DNA of nuclear or mitochondrial origin [18, 30, 39, 40], IFN-β signaling may also be triggered by cytosolic RNA as described in previous studies with X-irradiation [17, 18]. The RNA can be transcribed from cytosolic DNA or leak from mitochondria [41], but could also stem from reactivation of retroelements [42, 43] through radiation-induced decompaction of chromatin. Notably, a recent study showed a more pronounced IFN-β signaling after proton as compared to X-irradiation, which was caused by proton-induced derepression of transposable elements [44].

In conclusion, we have shown that ATR inhibition combined with low- or high-LET irradiation enhances type 1 IFN signaling in GBM cells. Furthermore, ATM inhibition can also enhance IFN signaling, as shown for one of the cell lines. The ATR and ATM inhibitors used in this study have been combined with classical radiotherapy in clinical trials for brain metastases (NCT02589522) and GBM (NCT03423628), respectively. GBM is typically highly radioresistant as well as invasive. Our results suggest that ATR or ATM inhibitors could increase tumor cell radiosensitivity and may also enhance radiation-induced immune signaling. The combination of these inhibitors with radiotherapy could thus likely facilitate eradication of the main tumor, and also help eliminating invasive cells outside the irradiation field by promoting antitumor immune effects. Likely, triple combinations of radiotherapy, radiosensitizing drug and immune checkpoint blockade might be most useful. In this approach the ATR/ATM inhibitor is used to increase tumor radiosensitivity and the antitumor immune effects of radiotherapy, while potential immunosuppressive effects, such as increased PD-L1 presentation, are counteracted by the immune checkpoint inhibitor.

## Author Contributions

Conceptualization (GER, MT, DIS, RGS); Formal analysis (GER, AEM, RGS); Funding acquisition (DIS, RGS); Investigation (GER, MT, AMS, AEM, AG); Methodology (NFJE, EM, FC, MT, GER); Project administration (DIS, RGS); Resources (DIS, FC, EM, RGS); Supervision (DIS, FC, RGS); Visualization (GER, AEM, MT, RGS);Writing - original draft (GER, RGS); and Writing - review & editing (all authors).

## Acknowledgements

This work was funded by Norway Romania Grant RO-NO-2019-0510, contract no. 41/2021grant. Irradiations with carbon ions were performed at Ganil, France, using beam time obtained under experiment number P1243-H of the iPAC 2020 call. The GANIL dosimetry team and beam operators are acknowledged for extensive support. We also acknowledge the operators at the Oslo Cyclotron Laboratory and members of the Malinen/Edin research group for excellent assistance with proton irradiation experiments.

