## Supplemental figures and table for "Interferon signaling is enhanced by ATR inhibition in glioblastoma cells irradiated with X-rays, protons or carbon ions"

**Supplementary figure legends**

**Suppl Fig S1:**

(A) Representative scatter plots from experiments used for quantifications in Figure 1C, showing signal intensity of γH2AX against DNA. For accurate comparison of signal intensity between samples, a barcoded X-irradiated sample (in red) was split between proton irradiated samples (in blue) before antibody staining. Upper panels show two similar experiments from T98G, in which the barcoded sample was irradiated with 6 Gy (top) and 12 Gy (bottom) of X-ray, showing comparable signal intensity to low- and high-LET protons, respectively. Similar results are shown for U-251 in the lower panels. (B) Left: Levels of γH2AX in T98G at 0.5 h after proton irradiation (shown in figure 1 C) displayed together with γH2AX levels at 0.5 h after various doses of X-irradiation ($n \geq 2$). Right: Remaining γH2AX signal at 24 h after irradiation relative to the signal at 0.5 h. Signal for 0 Gy has been subtracted before calculating the ratio.

**Suppl Fig S2:**

(A) DNA histogram obtained by flow cytometry analysis at 24 h after treatment with ATR and ATM inhibitors in combination with high-LET proton irradiation in T98G. (B) Quantification from cell cycle analysis performed on data as in A. Error bars: SEM ($n$ = 3, except for ATRi 100 nM ($n$ = 2).

**Suppl Fig S3:**

(A) Bar chart displaying the mean levels of secreted IFN-β from three experiments in U-251 cells co-treated with ATR or ATM inhibitors and X-irradiation as described in Figure 4. Individual experiments are shown as dotted lines. (B) Left: Quantification of pSTAT1 from immunoblot on proton irradiated T98G samples. Values are relative to total protein levels and normalized to the sample treated with 6 Gy high LET and 100 nM of ATR inhibitor, $n$ = 2. Right: Quantification of pSTAT1 from C-ion irradiated samples. Levels are relative to total protein and normalized to the sample treated with 4 Gy high LET and 100 nM of ATR inhibitor, $n$ = 4. (C) Correlation analysis between pSTAT1 and IFN-β secretion in X-irradiated samples of T98G. Signal intensity of pSTAT1 (relative to 6 Gy ATRi 250 nM) as measured by immunoblotting, is plotted against the level of secreted IFN-β measured in growth medium from the same sample relative to the amount of protein of the adherent cells. The IFN-β values have been normalized to the amount of protein of the adherent cells at time of harvest.

Supplementary figure 1

(A)

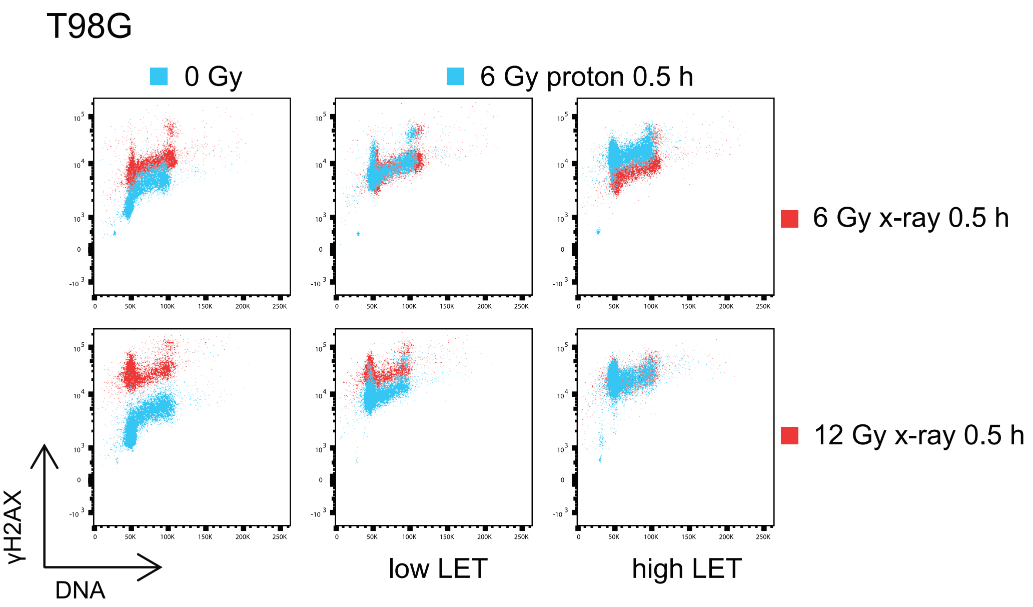


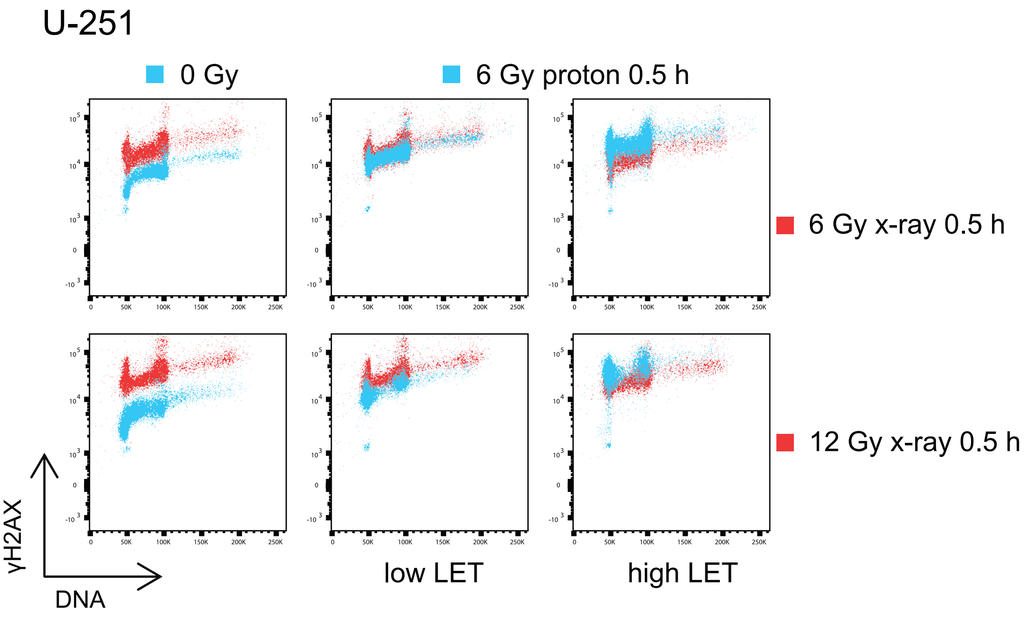


(B)

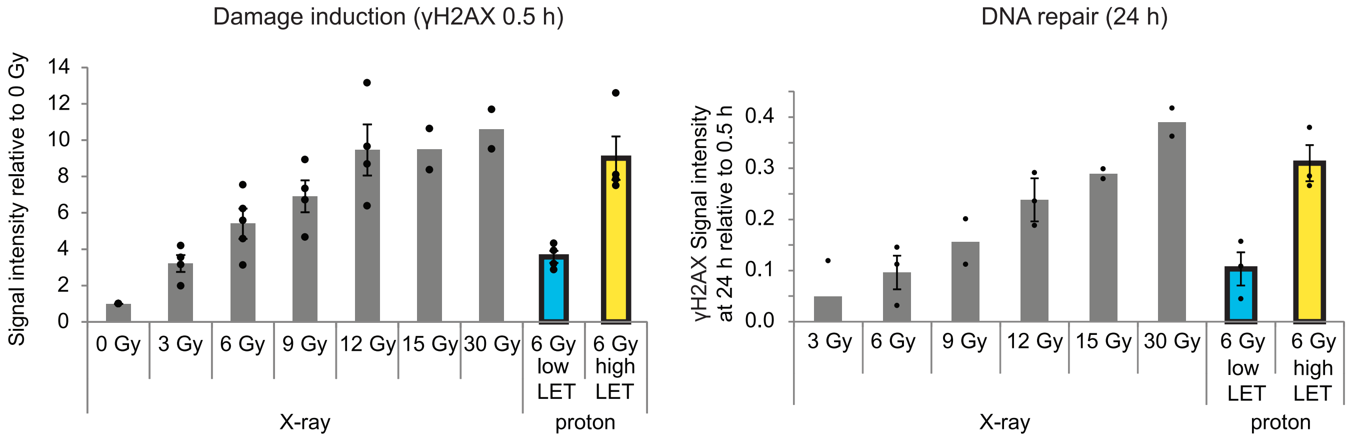

Supplementary figure 2

(A)

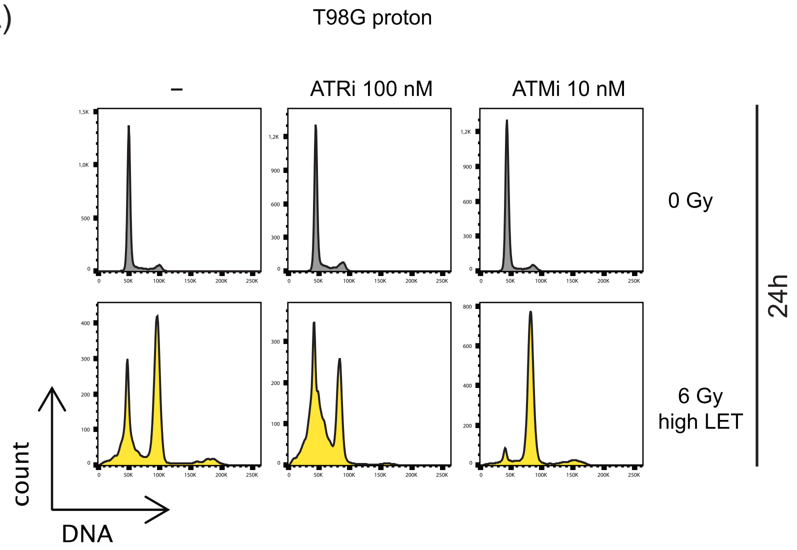


(B)

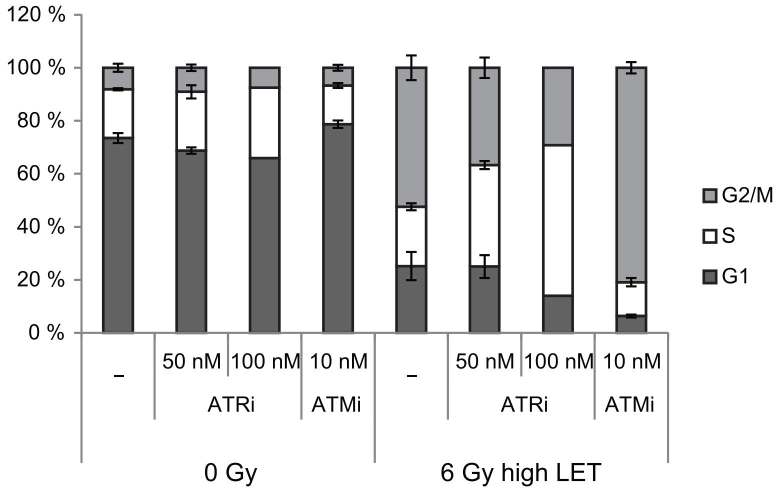

### Supplementary Figure 3

(A)

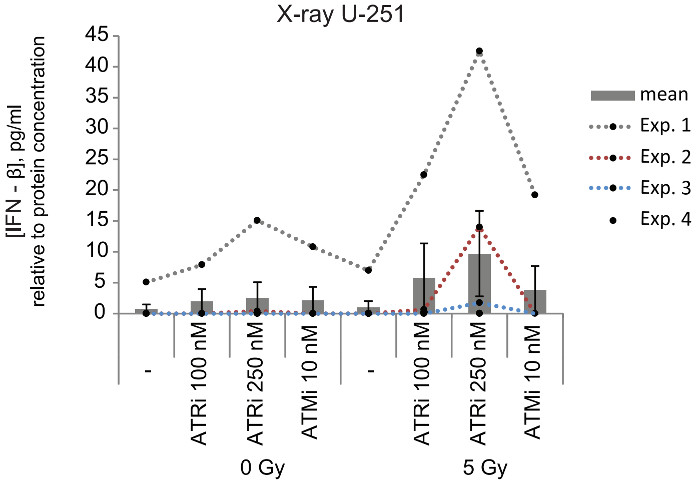


(B)

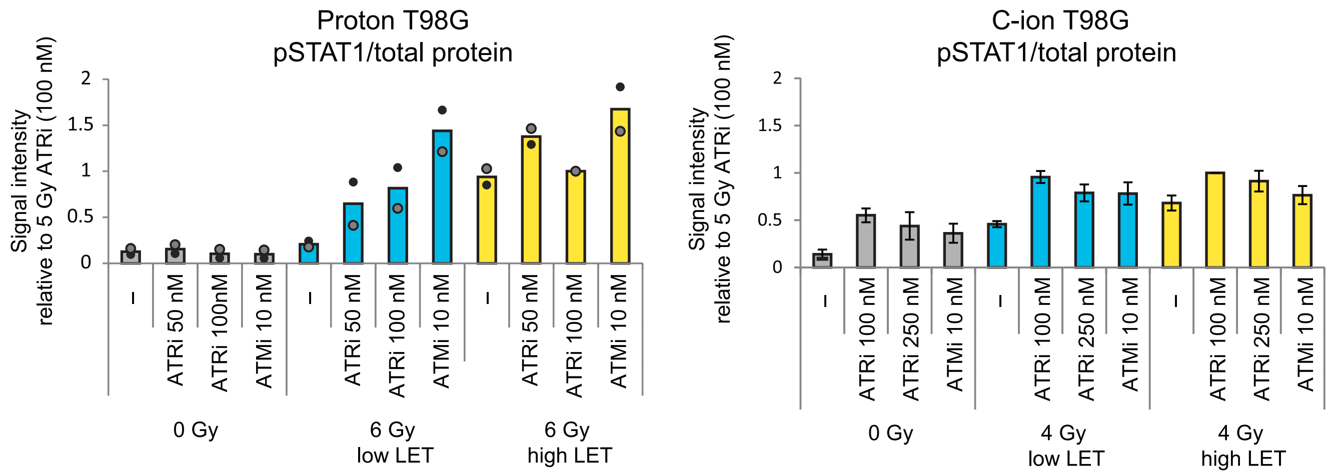


(C)

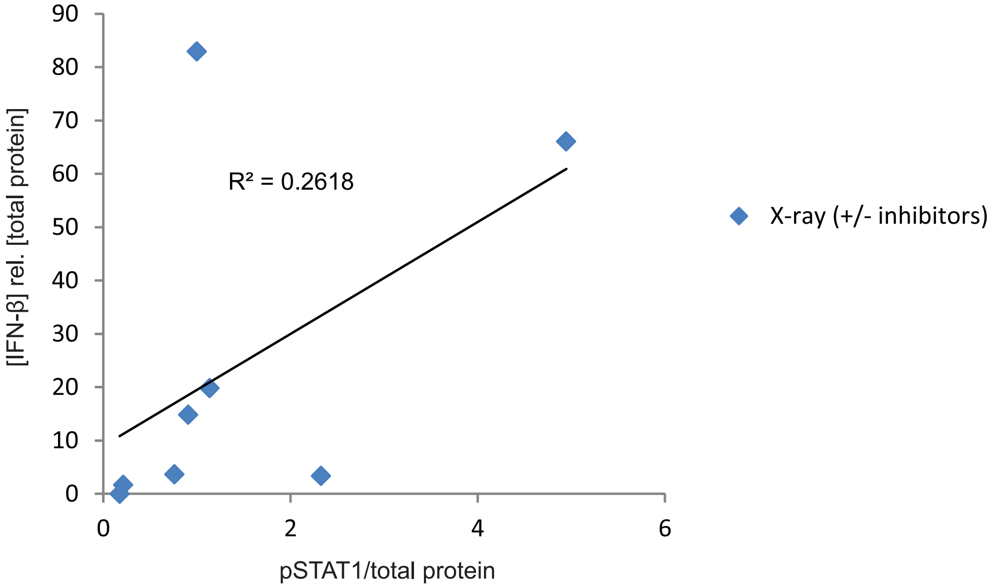

**Supplementary table 1** Antibodies for immunoblotting

| | Protein | Supplier | Cat. Nb. | Dilution |
|---|---|---|---|---|
| Primary antibodies | pATM S1981 | Cell Signaling Technology | #4526 | 1:1000 |
| | pCHK1 S345 | Cell Signaling Technology | #2348 | 1:400 |
| | pCHK1 S317 | Cell Signaling Technology | #2344 | 1:400 |
| | pSTAT1 Y701 | Cell Signaling Technology | #9167 | 1:1000 |
| | ATM (G-12) | Santa Cruz Biotechnology | #sc-377293 | 1:200 |
| | STAT1 | Santa Cruz Biotechnology | #sc-464 | 1:200 |
| | CHK1 (DCS310) | Invitrogen | #MA1-91087 | 1:200 |
| | γ-tubulin | Sigma-Aldrich | #T6557 | 1:2000 |
| Secondary antibodies | HRP goat anti-rabbit | Jackson ImmunoResearch | #111-035-144 | 1:10 000 |
| | HRP donkey anti-mouse | Jackson ImmunoResearch | #715-035-150 | 1:10 000 |

HRP: Horseradish peroxidase
